# The rVSV-EBOV vaccine provides limited cross-protection against Sudan virus in guinea pigs

**DOI:** 10.1101/2022.11.11.516195

**Authors:** Wenguang Cao, Shihua He, Guodong Liu, Helene Schulz, Karla Emeterio, Michael Chan, Kevin Tierney, Kim Azaransky, Geoff Soule, Nikesh Tailor, Abdjeleel Salawudeen, Rick Nichols, Joan Fusco, David Safronetz, Logan Banadyga

## Abstract

Recombinant vesicular stomatitis viruses (rVSVs) engineered to express heterologous viral glycoproteins have proven to be remarkably effective vaccines. Indeed, rVSV-EBOV, which expresses the Ebola virus (EBOV) glycoprotein, recently received clinical approval in the United States and Europe for its ability to prevent EBOV disease. Analogous rVSV vaccines expressing the glycoproteins of different human-pathogenic filoviruses have also been developed and shown to be effective in pre-clinical evaluations, yet these vaccine candidates have not progressed far beyond research laboratories. As the current outbreak of Sudan virus (SUDV) intensifies in Uganda, the need for proven countermeasures is rendered even more acute. Here we demonstrate that an rVSV-based vaccine expressing the SUDV glycoprotein (rVSV-SUDV) generates a potent humoral immune response that completely protects guinea pigs from SUDV disease and death. Although the cross-protection generated by rVSV vaccines for different filoviruses is thought to be limited, we wondered whether rVSV-EBOV might also provide protection against SUDV, which is closely related to EBOV. Surprisingly, nearly 60% of guinea pigs that were vaccinated with rVSV-EBOV and challenged with SUDV survived, suggesting that rVSV-EBOV offers limited protection against SUDV, at least in the guinea pig model. These results were confirmed by a back-challenge experiment in which animals that had been vaccinated with rVSV-EBOV and survived EBOV challenge were inoculated with SUDV and survived. Whether these data are applicable to the efficacy of rVSV-EBOV in humans is unknown, and they should therefore be interpreted cautiously. Nevertheless, this study confirms the potency of the rVSV-SUDV vaccine, and it highlights the potential for rVSV-EBOV to elicit a cross-protective immune response.

## Introduction

Filoviruses pose a significant threat to global public health *(1)*. Of the dozens of filoviruses that have so far been described, six of them are known to cause disease that is among the most severe viral illnesses observed in humans. Marburg virus (MARV) and Ravn virus (RAVV) both belong to the *Marburgvirus* genus and have been causing sporadic outbreaks mostly in Central Africa at least since the discovery of MARV in 1967. Moreover, MARV recently emerged for the first time in West Africa to cause a small outbreak in 2021 and another in 2022 *(2, 3)*. Ebola virus (EBOV), Bundibugyo virus (BDBV), Tai Forest virus (TAFV), and Sudan virus (SUDV) all belong to the *Ebolavirus* genus and have collectively caused numerous severe outbreaks throughout Africa since the first recorded appearance of EBOV and SUDV in 1976. Indeed, EBOV was responsible for the 2013-2016 West African epidemic that sickened more than 28,000 people and killed nearly half of them *(4)*. The West African EBOV epidemic was followed by several others, including a large outbreak in the Democratic Republic of the Congo in 2018-2020 that killed over 2000 people *(5)*. After EBOV and MARV, SUDV has been responsible for the most outbreaks, with four of the previous five occurring in Uganda, including the largest SUDV outbreak on record, which caused 425 cases and 224 deaths *(6)*. Uganda is currently in the midst of yet another SUDV outbreak, which has so far resulted in 136 confirmed cases and 53 deaths since the outbreak was declared on September 20, 2022 *(7)*.

The 2013-2016 West African EBOV epidemic stimulated remarkable progress in the pre-clinical and clinical development of filovirus countermeasures, although much of the groundwork for this rapid advancement had already been laid in the preceding decades of basic research *(8)*. In 2004, a recombinantly engineered vesicular stomatitis virus (rVSV) was first used as a vaccine to protect against disease caused by EBOV *(9)*. This vaccine was generated by removing the endogenous VSV glycoprotein (G) gene and replacing it with the gene for the EBOV glycoprotein (GP), which is responsible for virion entry and fusion. The resulting live, attenuated chimeric virus—known as rVSVΔG-ZEBOV-GP or, simply, rVSV-EBOV—expresses EBOV GP as the sole viral protein on the surface of the VSV virion. rVSV-EBOV has proven to be a remarkably safe, immunogenic, and effective vaccine, with years of research culminating in its recent licensure (Ervebo, Merck) in the United States and Europe for the prevention of EBOV disease *(10)*. The rVSV vaccine platform has also been used to develop a number of other experimental vaccines analogous to rVSV-EBOV *(11)*, including a SUDV vaccine (rVSV-SUDV) and a Lassa virus vaccine (rVSV-LASV).

Despite the recent advances made in countermeasure development, and despite a research pipeline full of promising experimental vaccines and therapeutics, there are still no approved treatments or prophylactics available for any filovirus other than EBOV. To date, only three SUDV-specific vaccine candidates have progressed to the point of Phase I clinical trials *(12)*, although the current outbreak in Uganda may stimulate additional trials. Two chimpanzee adenovirus-vectored vaccines—cAd3 expressing SUDV GP *(13)* and chAdOx1 expressing both EBOV and SUDV GP *(14, 15)*—are among the most promising vaccine candidates, although the Ad26.ZEBOV/MVA-BN-Filo prime-boost vaccine may also prove useful. Indeed, Ad26.ZEBOV/MVA-BN-Filo, which is already approved for use against EBOV (Zabdeno and Mvabea, Johnson & Johnson), may also confer protection against SUDV, thanks to the inclusion of SUDV GP in the MVA-BN-Filo boost *(16)*. Nevertheless, Ad26.ZEBOV/MVA-BN-Filo requires two injections, with Ad26.ZEBOV administered initially and MVA-BN-Filo administered as a boost. As a result, this vaccine may not be the best choice for outbreak response. rVSV-based vaccines, on the other hand, have demonstrated exceptional efficacy elicited relatively rapidly after only a single dose. Thus, in an effort to support the pre-clinical development of an rVSV-based SUDV vaccine, we used our guinea pig model of SUDV infection to evaluate our experimental rVSV-SUDV (also referred to as rVSVΔG-SUDV-GP), which expresses the GP of SUDV variant Boneface in place VSV G. Not surprisingly, our results confirm the effectiveness of this vaccine, which completely protected animals from SUDV disease and death, even when administered as a single injection at a relatively low dose. Not only do these data underscore the utility of rVSV-vectored vaccines against filoviruses, but they also lay a foundation for the future clinical development of this particular rVSV vaccine.

At the same time, however, we were also interested in knowing whether rVSV-EBOV was capable of providing cross-protection against SUDV challenge, since a readily available and clinically approved vaccine could have a meaningful benefit during an outbreak, even if it is only partially effective. Filoviruses are known to be serologically cross-reactive, and cross-reactivity may be higher among EBOV, SUDV, BDBV, and TAFV, which all belong to the *Ebolavirus* genus *(17–19)*. Indeed, the structural similarity between ebolavirus GPs is what has enabled the generation of broadly cross-reactive monoclonal antibodies, many of which were isolated from EBOV survivors but are capable of neutralizing multiple ebolaviruses *(20)*. However, while cross-protection may be possible in principle, limited reports with small numbers of nonhuman primates (NHPs) have suggested that rVSV-EBOV cannot protect against SUDV challenge *(21, 22)*. We therefore sought to re-evaluate the cross-protective efficacy of rVSV-EBOV in guinea pigs challenged with guinea pig-adapted (GPA) SUDV. Surprisingly, we observed that nearly 60% (8/14) of the guinea pigs that had been vaccinated with rVSV-EBOV survived challenge with SUDV, although most animals were not protected from disease. These results were confirmed by a back-challenge experiment in which rVSV-EBOV-immunized guinea pigs that had survived infection with GPA-EBOV were re-challenged with GPA-SUDV. Together, these data demonstrate that rVSV-EBOV is capable of eliciting an immune response that is cross-protective against SUDV in guinea pigs, but whether this phenomenon can be replicated in NHPs or humans remains to be determined.

## Results

### Immunization with rVSV-SUDV protects guinea pigs from lethal SUDV infection

To demonstrate the protective efficacy of rVSV-SUDV in the guinea pig model, two groups of 6 animals were immunized with either 2 × 10^5^ or 2 × 10^3^ PFU of rVSV-SUDV, and one group of 6 control animals was administered saline instead of vaccine. All animals were challenged with a uniformly lethal dose of GPA-SUDV on day 28 post-vaccination (Fig. S1A). All 6 control animals developed severe disease and succumbed to SUDV infection within 7-9 days, following significant weight loss and fever, defined as a body temperature greater than 39.5 °C for at least two consecutive days (Fig. 1A-C). Conversely, none of the vaccinated animals—regardless of vaccine dose—developed disease or succumbed to infection, with all animals consistently gaining weight and maintaining a normal body temperature throughout the study (Fig. 1A-C). A single vaccinated animal was lost during sampling on day 9 post-infection, but this was determined to be the result of a sampling accident. The animal itself was otherwise healthy and did not exhibit any signs of virus infection.

**Fig. 1.**
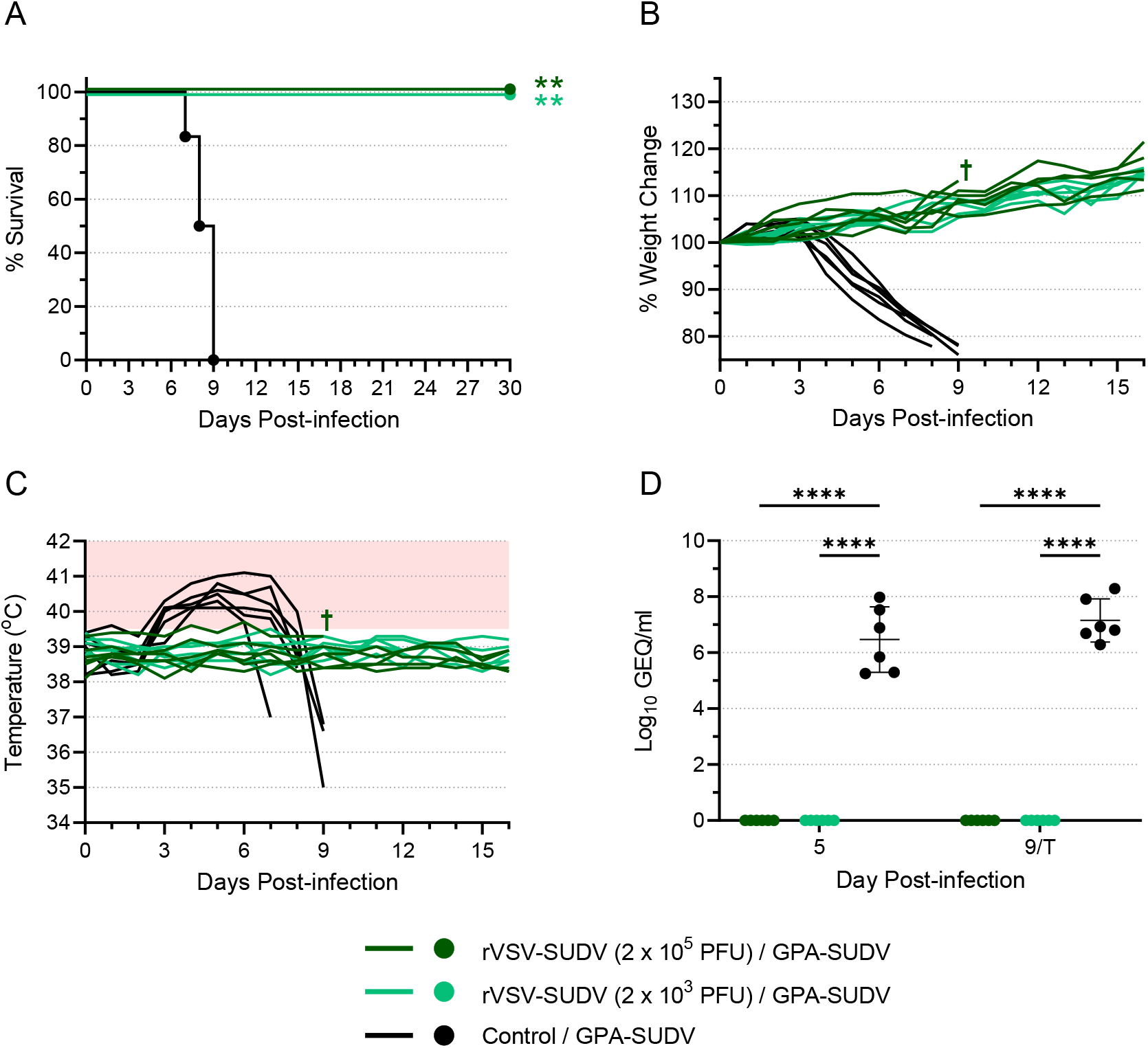
rVSV-SUDV protects against SUDV challenge in guinea pigs. Groups of 6 guinea pigs were vaccinated with rVSV-SUDV at a dose of either 2 × 10^5^ PFU or 2 × 10^3^ PFU. Control animals (n = 6) received an equivalent volume of saline. Twenty-eight days after vaccination, animals were challenged with 1000 LD_50_ of GPA-SUDV. Animals were monitored for survival (A), weight change (B), and body temperature (C). The area shaded pink in (C) highlights temperatures above 39.5 °C. Blood samples were obtained from each animal on day 5 post-infection and either day 9 post-infection or at the terminal time point (T) if it occurred before day 9. Samples were assessed for levels of virus RNA via RT-qPCR, and data are presented as Log_10_ genome equivalents (GEQ) per milliliter for each animal, with means and standard deviations indicated (D). A single animal (†) that was vaccinated with 2 × 10^5^ PFU rVSV-SUDV died during sampling on day 9; this animal did not exhibit signs of disease and is considered a survivor. **, p ≤ 0.01; ****, p ≤ 0.0001.

Levels of virus RNA in the blood were assessed by RT-qPCR for all animals on day 5 post-infection and again on either day 9 or the terminal time point (prior to euthanasia), if it occurred before day 9 (Fig. 1D). Remarkably, none of the immunized animals exhibited detectable levels of SUDV RNA, suggesting that viremia was prevented by immunization with rVSV-SUDV. In contrast, the control animals exhibited significantly higher levels of virus RNA on both day 5 and the terminal time point, reflective of the severe disease they experienced.

Animals were likely protected from SUDV disease by the potent humoral immune response elicited by vaccination. Indeed, an ELISA revealed low levels of SUDV GP-specific IgG as early as 7 days post-vaccination, with endpoint titers in all animals increasing by day 14 and remaining high until the time of SUDV challenge (Fig. 2A). By day 28 post-vaccination, the geometric mean endpoint titers for all vaccinated animals were around 4.5 Log_10_. As expected, the control animals did not mount an IgG response to SUDV GP. Interestingly, immunization with rVSV-SUDV did not elicit appreciable levels of EBOV GP-specific IgG in most animals (Fig. 2B), suggesting the lack of a cross-protective humoral immune response, at least given the vaccine dose levels and administration regimen used here. Regardless, these data confirm the effectiveness of rVSV-SUDV immunization against GPA-SUDV challenge, demonstrating 100% protection from morbidity and mortality for the first time in the guinea pig model.

**Fig. 2.**
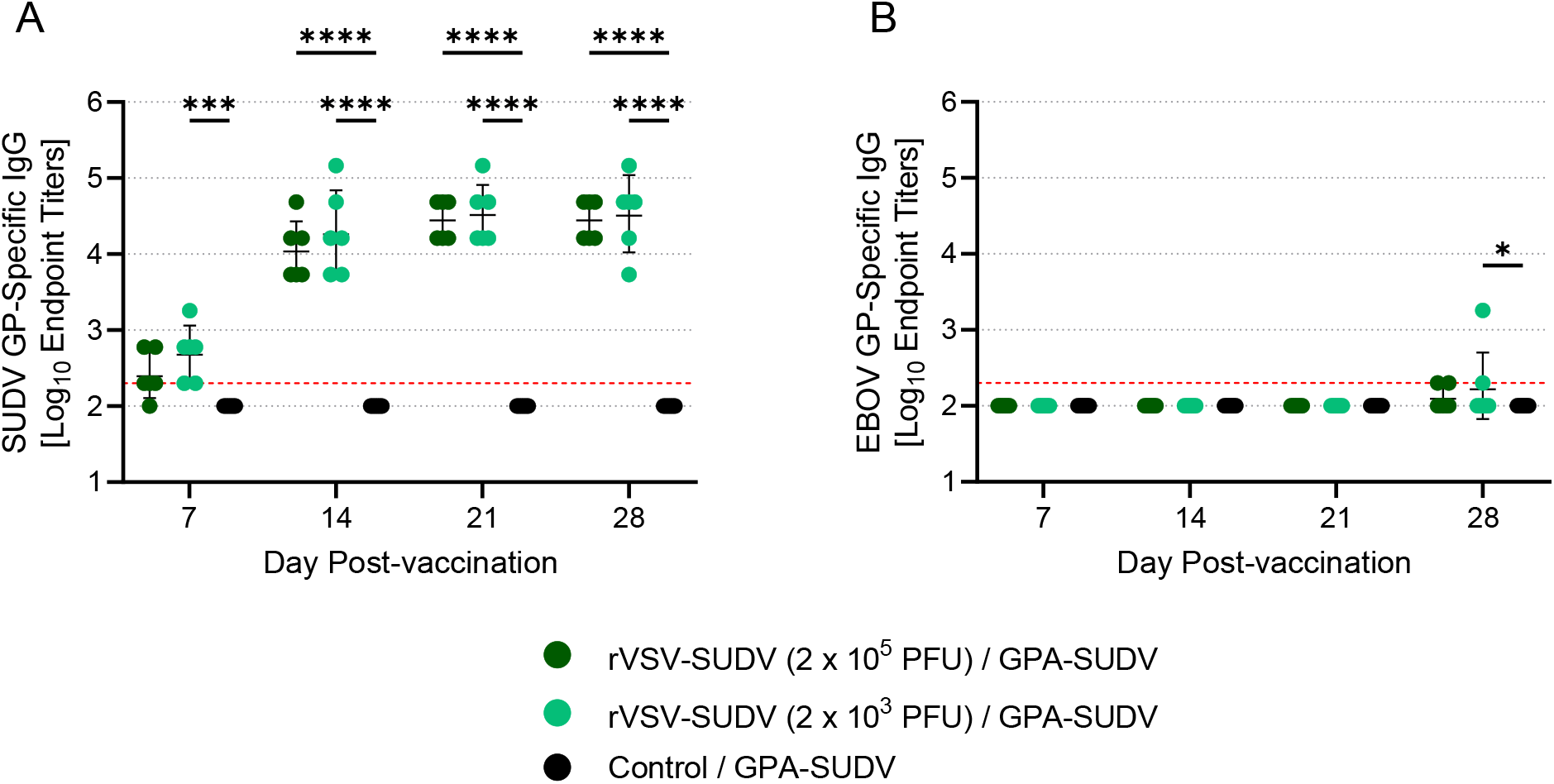
rVSV-SUDV elicits a robust humoral immune response. Serum samples were obtained from all animals 28 days after vaccination with rVSV-SUDV and prior to challenge with GPA-SUDV. Samples were assessed for levels of SUDV GP-specific IgG (A) or EBOV GP-specific IgG (B) via ELISA. Data are presented as Log_10_ endpoint titers for each animal, with the geometric means and standard deviations indicated. The lower limit of detection is indicated with a red dashed line. *, p ≤ 0.05; ***, p ≤ 0.001; ****, p ≤ 0.0001.

### Immunization with rVSV-EBOV provides limited cross-protection from lethal SUDV infection

To determine whether immunization with rVSV-EBOV could offer heterologous protection against SUDV challenge in the guinea pig model, 20 guinea pigs were immunized with 1 × 10^6^ PFU of rVSV-EBOV and, 28 days later, 14 animals were challenged with GPA-SUDV while the remaining 6 were challenged with GPA-EBOV (Fig. S1C, E). Ten guinea pigs immunized with rVSV-LASV were used as controls, with 5 animals challenged with GPA-SUDV and the other 5 challenged with GPA-EBOV.

Unsurprisingly, all animals that were vaccinated with rVSV-EBOV and challenged with GPA-EBOV survived infection (Fig. 3A), showing no signs of disease, weight loss, or fever throughout the study (Fig. 3B, C). Although EBOV RNA was detected in the blood of two vaccinated animals on day 5 post-infection, no virus RNA was detectable by day 9 (Fig. 3D). In contrast, all animals vaccinated with rVSV-LASV succumbed to EBOV infection by day 8 post-infection, exhibiting dramatic weight loss, a spike in body temperatures, and very high levels of virus RNA in blood samples collected on day 5 and at the terminal time point (Fig. 3A-D). These data confirm the outstanding protective efficacy of rVSV-EBOV against EBOV challenge.

**Fig. 3.**
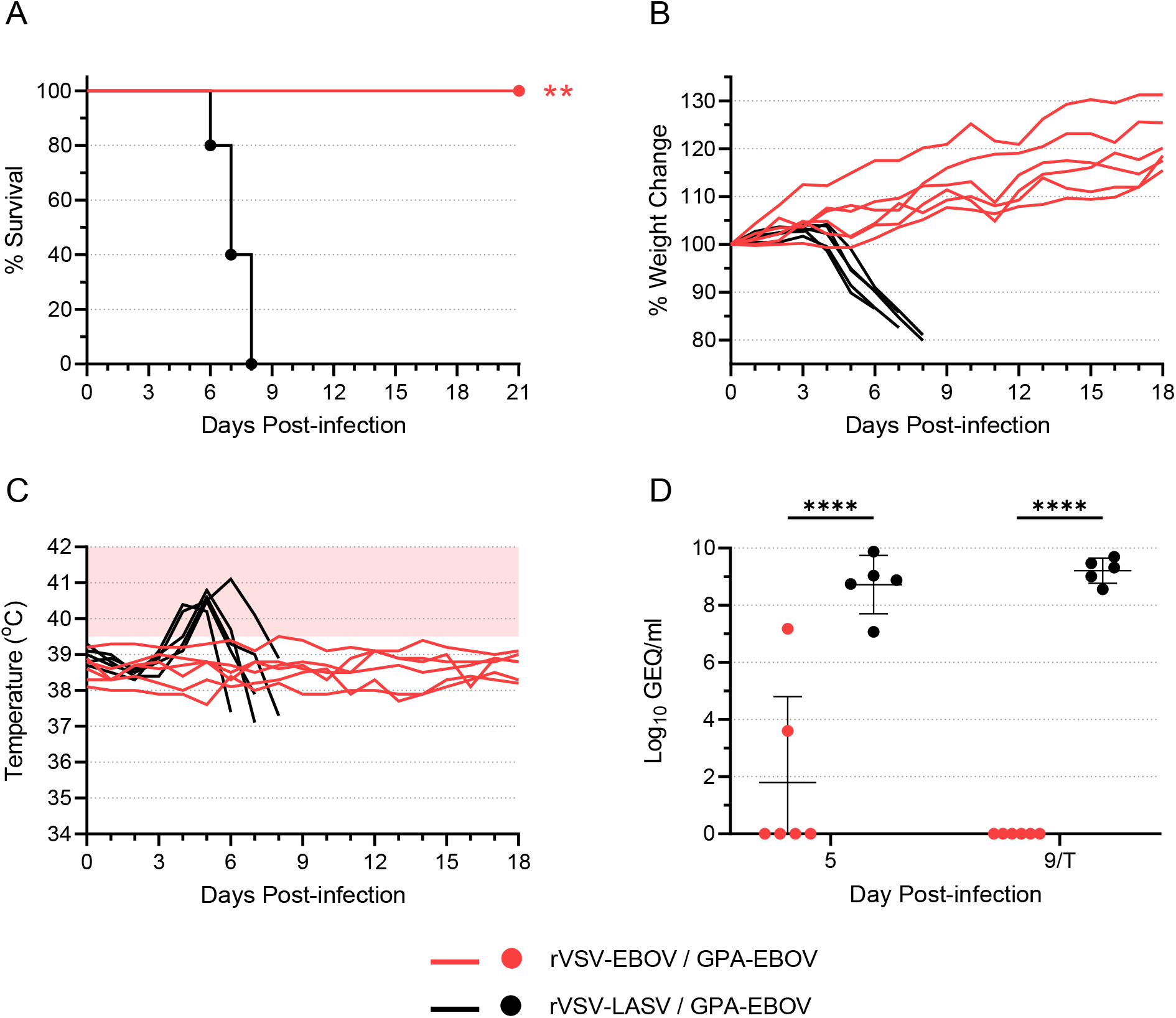
rVSV-EBOV protects against EBOV challenge in guinea pigs. Guinea pigs were vaccinated with rVSV-EBOV (n = 6) or rVSV-LASV (n = 5) at a dose of 1 × 10^6^ PFU. Twenty-eight days after vaccination, animals were challenged with 1000 LD_50_ of GPA-EBOV. Animals were monitored for survival (A), weight change (B), and body temperature (C). The area shaded pink in (C) highlights temperatures above 39.5 °C. Blood samples were obtained from each animal on day 5 post-infection and either day 9 post-infection or at the terminal time point (T) if it occurred before day 9. Samples were assessed for levels of virus RNA via RT-qPCR, and data are presented as Log_10_ genome equivalents (GEQ) per milliliter for each animal, with means and standard deviations indicated (D). **, p ≤ 0.01; ****, p ≤ 0.0001.

Remarkably, of the 14 guinea pigs that were vaccinated with rVSV-EBOV and challenged with GPA-SUDV, 8 animals survived (Fig. 4A; Fig. S2). Of the surviving animals, 5 showed signs of moderate disease, with animals losing between 8-15% of their body weight and most exhibiting a mild to moderate fever (Fig. 4B, C; Fig. S2). The remaining 3 survivors lost very little weight (<5%, if any) and showed no outward signs of disease, although two of them did exhibit a fever (Fig. 4B, C; Fig. S2). In contrast, the 6 rVSV-EBOV-vaccinated animals that succumbed to SUDV all exhibited severe signs of SUDV disease, with significant weight loss and pronounced fevers (Fig. 4A-C; Fig. S2). Likewise, the control animals that were vaccinated with rVSV-LASV and challenged with GPA-SUDV also exhibited severe signs of disease, succumbing between days 7 and 11 post-infection following significant weight loss and fever (Fig. 4A-C).

**Fig. 4.**
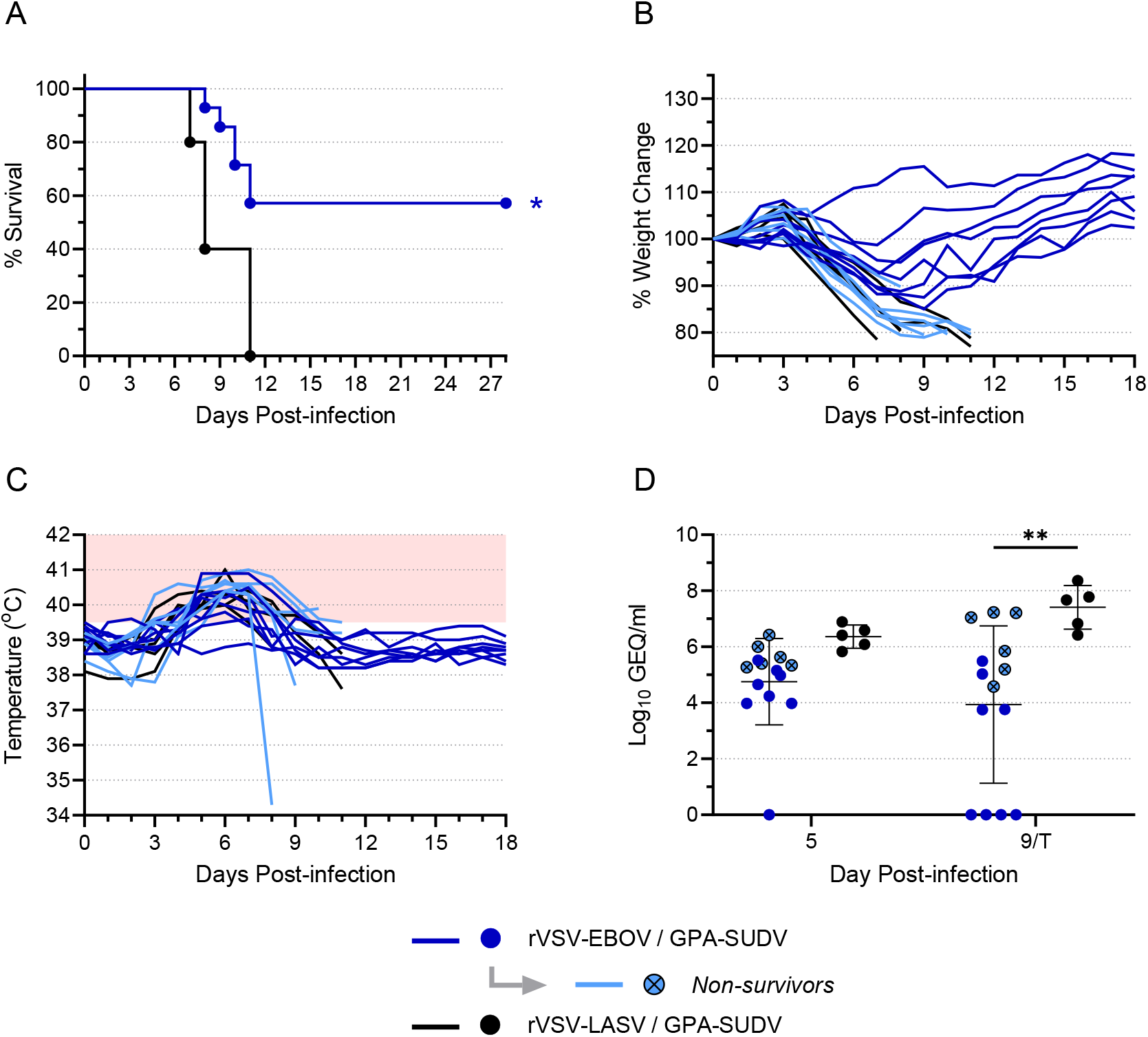
rVSV-EBOV provides limited cross-protection against SUDV challenge in guinea pigs. Guinea pigs were vaccinated with rVSV-EBOV (n = 14) or rVSV-LASV (n = 5) at a dose of 1 × 10^6^ PFU. Twenty-eight days after vaccination, animals were challenged with 1000 LD_50_ of GPA-SUDV. Animals were monitored for survival (A), weight change (B), and body temperature (C). The area shaded pink in (C) highlights temperatures above 39.5 °C. Blood samples were obtained from each animal on day 5 post-infection and either day 9 post-infection or at the terminal time point (T) if it occurred before day 9. Samples were assessed for levels of virus RNA via RT-qPCR, and data are presented as Log_10_ genome equivalents (GEQ) per milliliter for each animal, with means and standard deviations indicated (D). Data from animals that were vaccinated with rVSV-EBOV but did not survive challenge with GPA-SUDV are highlighted in light blue and indicated with an “x” on the symbol. *, p ≤ 0.05; **, p ≤ 0.01.

At day 5 post-infection, SUDV RNA was detected in most animals—regardless of vaccination (Fig. 4D). All but one of the 8 surviving animals showed moderate levels of virus RNA, while the 6 non-survivors all exhibited RNA levels that on average, trended higher than that of the survivors, although the difference was not statistically significant (Fig. S3). The rVSV-LASV-vaccinated animals had slightly higher levels of virus RNA, but this was also not statistically different compared to the rVSV-EBOV-vaccinated animals. At day 9 post-infection, four of the surviving animals had no detectable SUDV RNA, while the other four survivors showed moderate levels of RNA (Fig. 4D). All non-surviving animals had significantly higher levels of RNA than the survivors (Fig. S3). Although the overall difference in RNA levels between the rVSV-EBOV-and rVSV-LASV-vaccinated animals was statistically significant (Fig. 4D), the difference between the non-survivors in each vaccine group was not (Fig. S3). Interestingly, unlike the animals vaccinated with rVSV-SUDV, which did not exhibit an IgG response against EBOV GP, all rVSV-EBOV-vaccinated animals exhibited a heterologous IgG response against SUDV GP prior to challenge, albeit to a lesser degree than EBOV GP (Fig. 5).

**Fig. 5.**
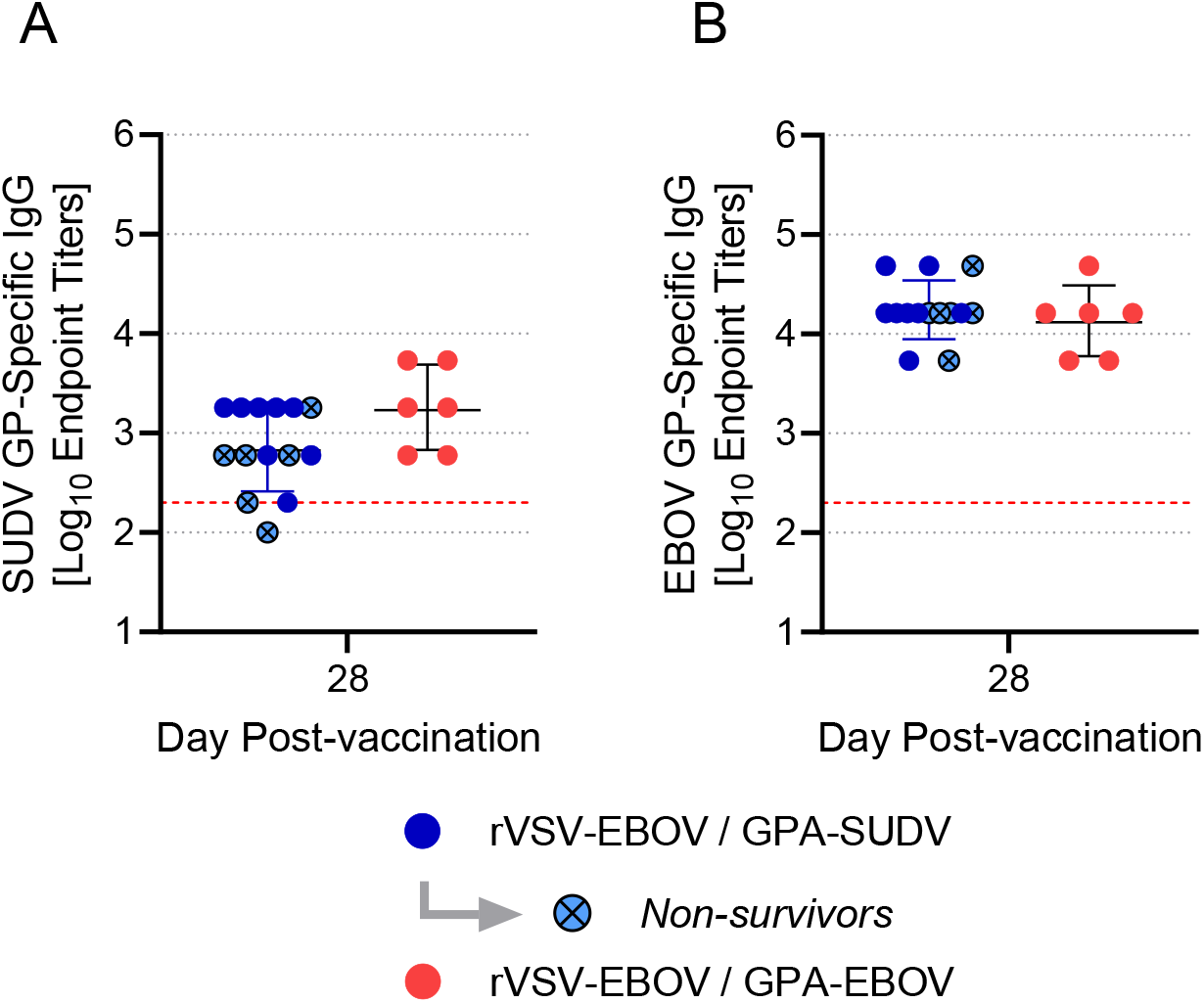
rVSV-EBOV elicits a cross-reactive humoral immune response. Serum samples were obtained from all animals 28 days after vaccination with rVSV-EBOV and prior to challenge with GPA-SUDV. Samples were assessed for levels of SUDV GP-specific IgG (A) or EBOV GP-specific IgG (B) via ELISA. Data are presented as Log_10_ endpoint titers for each animal, with the geometric means and standard deviations indicated. The lower limit of detection is indicated with a red dashed line. Data from animals that were vaccinated with rVSV-EBOV but did not survive challenge with GPA-SUDV are highlighted in light blue and indicated with an “x” on the symbol.

### Back-challenge with EBOV but not SUDV results in lethal disease

To further investigate the degree of cross-protection elicited by vaccination and infection, we performed two back-challenge experiments (Fig. S1B, D). The 11 guinea pigs that were vaccinated with rVSV-SUDV were back-challenged with GPA-EBOV 30 days after they were originally challenged with GPA-SUDV (Fig. 1), and the 6 guinea pigs that were vaccinated with rVSV-EBOV were back-challenged with GPA-SUDV 21 days after they were originally challenged with GPA-EBOV (Fig. 3). All animals that were back-challenged with GPA-SUDV survived (Fig. S4A), and the majority showed no signs of illness. One of the 6 animals did exhibit weight loss and a mild fever (Fig. S4B, C), coinciding with moderate levels of virus RNA in the blood at day 5 (Fig. S4D), but this animal recovered completely. In contrast, 8 of the 11 animals (∼73%) that were back-challenged with GPA-EBOV died after exhibiting severe disease, with the survival curves showing no significant difference from that of control animals (Fig. S5A-C). All non-surviving animals exhibited high levels of virus RNA in the blood (Fig. S5D). The three surviving animals remained disease-free throughout the experiment, and showed no detectable virus RNA at either day 5 or day 10 post-infection (Fig. S5A-D). These data further demonstrate that rVSV-EBOV vaccination (followed by EBOV challenge) confers cross-protection against SUDV; however, they also suggest that the converse scenario does not hold true. The majority of animals that were vaccinated with rVSV-SUDV and survived SUDV challenge were not able to overcome infection with GPA-EBOV, suggesting a lack of a cross-protective immune response.

## Discussion

The results presented here demonstrate that it is possible, in principle, to achieve cross-protection against challenge with SUDV following rVSV-EBOV immunization—at least in guinea pigs. Nearly 60% of the guinea pigs that were immunized with 1 × 10^6^ PFU of rVSV-EBOV survived challenge with SUDV 28 days later. Prior to challenge, almost all of these animals exhibited moderate levels of SUDV GP-specific IgG, in addition to high levels of EBOV GP-specific IgG. Since the humoral immune response is key to the protection offered by rVSV-EBOV *(23)*, it seems reasonable to assume that it was also responsible for providing partial protection against heterologous challenge with SUDV, although we cannot rule out the contribution of other arms of the immune response. Moreover, 100% of the guinea pigs that survived GPA-EBOV infection after rVSV-EBOV immunization also survived GPA-SUDV back-challenge. Despite the small number of animals, this result may suggest that the cross-protective immune response was boosted following GPA-EBOV infection. Although these data challenge the notion that rVSV-based filovirus vaccines cannot confer cross protection, the results must not be over-interpreted. These experiments were performed using a guinea pig-adapted virus in guinea pigs, which reliably recapitulate many of the hallmarks of filovirus disease and are used routinely to study EBOV, SUDV, and other filoviruses *(24)*. However, guinea pigs are not humans, and results in guinea pigs may not necessarily predict what will happen in humans. The cross-protective efficacy of rVSV-EBOV should ideally be confirmed in the NHP model, which is considered the gold-standard animal model for filoviruses *(25)*. While encouraging, the results presented here should not be used alone to make decisions about the clinical applications of the licensed rVSV-EBOV vaccine in the context of related filovirus outbreaks. Rather, these results should be used to stimulate further research into the immune responses elicited by filovirus vaccines and the potential for cross-protective efficacy.

The ability of rVSV-based filovirus vaccines to elicit meaningful protection against heterologous viruses is difficult to sort out based on the handful of previously published reports. Moreover, the degree of cross-protection is likely influenced by a number of variables, including the animal model, the magnitude and duration of the immune response, the vaccine virus and dose, and the challenge virus. For instance, Marzi et al. demonstrated robust cross-protection conferred by rVSV vaccines expressing TAFV, SUDV, or Reston virus GP against mouse-adapted EBOV in mice, yet the cross-protection was essentially lost when the same experiment was performed using GPA-EBOV in guinea pigs *(26)*.

Similarly, the results from our back-challenge experiment showed that vaccination with rVSV-SUDV (followed by infection with GPA-SUDV) was not capable of protecting against GPA-EBOV. The reasons for this lack of cross-protection are unclear—especially in light of the cross-protection offered by rVSV-EBOV—but they could be related to differences in the quality of immune response elicited by each vaccine and/or differences in virus pathogenicity. In cynomolgus macaques, 2 × 10^7^ PFU of rVSV-EBOV provided partial protection following BDBV challenge, but rVSV-TAFV did not *(27)*. A similar experiment demonstrated that a prime with 1 × 10^7^ PFU rVSV-SUDV followed by a boost with the same dose of rVSV-EBOV could offer partial protection against BDBV challenge, but a blended vaccine consisting of rVSV-SUDV and rVSV-EBOV was not effective *(28)*. Conversely, a blended vaccine consisting of rVSV-EBOV, rVSV-SUDV, and rVSV-MARV conferred cross-protection against challenge with TAFV, but a single animal immunized with 3 × 10^7^ PFU rVSV-EBOV and then challenged 28 days later with SUDV did not survive *(22)*. Likewise, Jones et al. reported that cynomolgus macaques that had survived EBOV infection following rVSV-EBOV vaccination (1 × 10^7^ PFU) were not protected from subsequent challenge with SUDV variant Gulu, with 3 of the 4 animals dying *(21)*. Because SUDV is not uniformly lethal in cynomolgus macaques, it is difficult to know whether the lone surviving animal in this case was protected by rVSV-EBOV vaccination or would have survived SUDV infection regardless. Interestingly, a recent report has demonstrated that immunization of cynomolgus macaques with 5 × 10^7^ PFU of rVSV-EBOV followed by infection with EBOV results in a SUDV GP-specific IgG response that persists, albeit at low levels, up to 290 days post-vaccination *(29)*. However, following a control immunization with rVSV-MARV, these animals were not protected from subsequent challenge with SUDV, with 4 of 5 succumbing to disease. Collectively, these data suggest that rVSV-EBOV cannot be relied upon to generate an immune response that is sufficiently cross-protective against SUDV. Why, then, did we observe partial cross-protection in guinea pigs immunized with rVSV-EBOV? While the guinea pig model, in general, is considered to be more stringent and predictive than the mouse model *(30)*, it is likely still less stringent than the NHP model and may offer a lower bar for protection than is required in monkeys. Additionally, guinea pigs are naturally resistant to infection with wild type filoviruses, including SUDV *(31)*. Our model therefore relies on a guinea pig-adapted strain of SUDV *(32)*, which contains a number of point mutations that confer virulence in guinea pigs but may not completely recapitulate the pathogenic processes that occur in NHPs and humans. Finally, the duration between vaccination and challenge likely also affects the outcome. As vaccine-elicited antibody titers are expected to decrease over time, it is unclear to what degree rVSV-EBOV would continue to offer protection against SUDV beyond 28 days post-vaccination. Clearly, there are many questions left to answer regarding the cross-protective potential of rVSV-EBOV, underscoring the necessity for follow-up experiments, including evaluation in NHPs.

Whether or not rVSV-EBOV is capable of eliciting meaningful cross-protection against SUDV in humans, it is worth remembering that rVSV-SUDV is already known to offer robust protection against SUDV. Our data demonstrate that rVSV-SUDV elicits a uniformly high SUDV GP-specific antibody response in guinea pigs as early as 14 days after a single administration and even when given at a relatively low dose of 2 × 10^3^ PFU. These data confirm that the rVSV-SUDV vaccine is highly effective at preventing Sudan virus disease. Additionally, Marzi et al. have recently demonstrated the effectiveness of rVSV-SUDV in NHPs *(29)*, and a past report showed that the vaccine also works as a post-exposure treatment *(33)*. Given the success of the rVSV-EBOV vaccine, and abundant data demonstrating the efficacy of analogous VSV vaccines for other filoviruses, it is long past time for rVSV-SUDV to be advanced to clinical trials.

## Materials and Methods

### Animal ethics and biosafety statement

All animal experiment protocols were reviewed and approved by the Animal Care Committee at the Canadian Science Centre for Human and Animal Health (CSCHAH), Winnipeg, Manitoba, in accordance with guidelines from the Canadian Council on Animal Care (CCAC). All staff working on animal experiments completed education and training programs according to the standard protocols appropriate for this level of biosafety. All work with infectious SUDV and EBOV was performed in the containment level (CL)-4 laboratories at the CSCHAH in accordance with standard operating protocols.

### Viruses

Guinea pig-adapted Sudan virus variant Boneface (GPA-SUDV; Sudan virus/NML/C.porcellus-lab/SSD/1976/Nzara-Boneface-GP; Genbank accession number KT750754.1) and guinea pig-adapted Ebola virus variant Mayinga (GPA-EBOV) were generated as previously described *(32, 34)*. Recombinant vesicular stomatitis virus (rVSV)-based vaccines expressing SUDV GP, EBOV GP, or LASV G in place of VSV G were generated as previously described *(9)*.

### rVSV-SUDV efficacy against GPA-SUDV in guinea pigs

To evaluate the protective efficacy of rVSV-SUDV immunization against GPA-SUDV challenge in female Hartley guinea pigs (Charles River Laboratories), groups of 6 animals each were immunized via the intramuscular route with either 2 × 10^5^ PFU or 2 × 10^3^ PFU of rVSV-SUDV (Fig. S1A). A control group of 6 animals received an equivalent volume of 0.9% saline. Twenty-eight (28) days post-vaccination, all animals were inoculated with 1000 times the median lethal dose (LD_50_) of GPA-SUDV via intraperitoneal (IP) injection. Animals were monitored for disease and survival up to 30 days post-infection. Weights were recorded daily for all animals up to day 16, as were body temperatures (as measured via subcutaneously implanted transponders). EDTA blood and/or serum samples were obtained from all animals prior to vaccination, on day 5 post-infection, and on either day 9 post-infection or at the animal’s terminal time point if it occurred before day 9.

On day 30 post-infection, all immunized animals (n = 11) were back-challenged with 1000 LD_50_ of GPA-EBOV via IP injection (Fig. S1B). EDTA blood and/or serum samples were obtained prior to back-challenge, on day 5 post-infection, and on either day 10 post-infection or at each animal’s terminal time point if it occurred before day 10. Animals were monitored for disease and survival up to 14 days post-infection with weights and temperatures recorded daily.

### rVSV-EBOV efficacy against GPA-SUDV in guinea pigs

To evaluate the cross-protective efficacy of rVSV-EBOV immunization against GPA-SUDV challenge in female Hartley guinea pigs (Charles River Laboratories), 20 animals were immunized via the IP route with 1 × 10^6^ PFU of rVSV-EBOV (Fig. S1C, E). A control group of 10 animals were immunized via the IP route with 1 × 10^6^ PFU of rVSV-LASV. Twenty-eight (28) days post-vaccination, 14 of the animals that had been immunized with rVSV-EBOV were challenged with 1000 LD_50_ of GPA-SUDV via the IP route, while the remaining 6 were challenged with 1000 LD_50_ of GPA-EBOV. At the same time point, 5 of the control animals (immunized with rVSV-LASV) were challenged with 1000 LD_50_ of GPA-SUDV, and the other 5 were challenged with 1000 LD_50_ of GPA-EBOV. Animals challenged with GPA-EBOV were monitored for disease and survival up to day 21 post-infection, while animals challenged with GPA-SUDV were monitored for disease and survival up to day 28. Weights were recorded daily for all animals up to day 18, as were body temperatures (as measured via subcutaneously implanted transponders). EDTA blood and/or serum samples were obtained from all animals prior to vaccination, on day 5 post-infection, and on either day 9 post-infection or at the animal’s terminal time point, if it occurred before day 9.

On day 21 post-infection, all animals (n = 6) that had been immunized with rVSV-EBOV and challenged with GPA-EBOV were back-challenged with 1000 LD_50_ of GPA-SUDV via IP injection (Fig. S1D). EDTA blood and/or serum samples were obtained prior to back-challenge and on days 5 and 10 post-infection. Animals were monitored for disease and survival up to 14 days post-infection with weights and temperatures recorded daily.

### Virus RNA quantification

EDTA blood samples were inactivated using Buffer AVL (Qiagen) and ethanol, according to manufacturer’s instructions. Viral RNA was extracted from these samples using the KingFisher Viral NA Kit (Thermo Fisher) on the KingFisher Apex per the manufacturer’s protocol. GPA-SUDV and GPA-EBOV RNA levels were determined by reverse transcription quantitative PCR (RT-qPCR) using the TaqPath 1-Step Multiplex Master Mix (Thermo Fisher) on the Applied Biosystems QuantStudio 3, along with the SUDV-specific primers (forward, 5′-CAGAAGACAATGCAGCCAGA-3′; reverse, 5′-TTGAGGAATATCCCACAGGC-3′; probe, 5′-6-FAM-CTGCTAGCT/Zen/TGGCCAAAGTCACAAG-IABkFQ-3′) or EBOV-specific primers (forward, 5′-CAGCCAGCAATTTCTTCCAT-3′; reverse, 5′-TTTCGGTTGCTGTTTCTGTG-3′; probe 5′-6-FAM-ATCATTGGCGTACTGGAGGAGCAG-IABkFQ-3′).

Cycling conditions were as follows: 25°C for 2 min, 53°C for 10 min and 95°C for 2 min, followed by 40 cycles of 95°C for 3 s and 60°C for 30 s. Standard curves were generated from plasmids encoding SUDV L or EBOV L and were used to convert the cycle threshold (Ct) values to genome equivalents per milliliter (GEQ/mL).

### IgG ELISAs

SUDV GP-specific IgG and EBOV-specific IgG levels were quantified in serum samples by indirect ELISA. Half-area high-binding 96-well assay plates (Corning) were coated using transmembrane domain-deleted SUDV GP or EBOV GP proteins (IBT Bioservices) prepared in pH 9.5 carbonate buffer in a 30-ul volume (1 ug/ml) at 4 °C for overnight. On the day of the experiment, after removing the coating solution, plates were incubated with 100 ul of 5% skim milk (BD Biosciences) prepared in 0.1% Tween-20 in PBS for 3 h at 37 °C. Serial dilutions of serum samples prepared in 2% milk were then applied to the plates (30 ul/well) and allowed to incubate at 4 °C overnight. Following 4 washes with 0.1% Tween-20/PBS, the plates were incubated for 1 h at 37 °C with an HRP-conjugated goat anti-guinea pig IgG(H+L) secondary antibody (KPL) diluted 1:5000 in 2% milk, followed by 4 washes with 0.1% Tween-20/PBS. The plates were then incubated in a TMB solution (Life Technologies) for ∼30 min in darkness before optical density (OD) signals were measured at 650 nm using a Synergy HTX plate reader (Biotek). Endpoint dilution titers were calculated by determining the highest dilution that gave an average OD 650 reading greater than or equal to the cut-off OD value, which was set as the mean OD value for pre-vaccine serum samples plus three times the standard deviation. When the endpoint titer was determined to lie below the lower limit of detection, an arbitrary value of 1:100 was assigned.

### Statistical analyses

GraphPad Prism version 9 was used to perform all statistical tests and generate all graphs. The Kaplan-Meier survival curves were compared using the Log-Rank test with the Bonferroni correction for multiple comparisons (where necessary). The two-way ANOVA test, along with Tukey’s multiple comparison test, was used to compare the means in Figures 1D, 2, 3D, 4D, S3, S4D, and S5D. An unpaired t-test was used to compare the means in Figure 5. Statistically significant differences are indicated with asterisks, where a P-value less than or equal to 0.05 was marked with one asterisk (*), less than or equal to 0.01 was marked with two asterisks (**), less than or equal to 0.001 was marked with three asterisks (***), and less than or equal to 0.0001 was marked with four asterisks (****).

## Author Contributions

L.B., S.H., D.S., R.N., and J. F. conceived the experiments described in the manuscript. W.C., S.H., G.L., H.S., K.E., M.C., K.T., K.A., G.S., N.T., and A.S. performed the experiments. W.C., S.H., G.L., D.S, and L.B. analyzed the data. L.B. performed the statistical analyses, created the figures, and wrote the first draft of the manuscript. W.C., G.L., R.N., J.F., D.S., and L.B. contributed to revising and editing the manuscript. All authors have read the manuscript and approve its content.

## Supplementary Materials

**Fig. S1.**
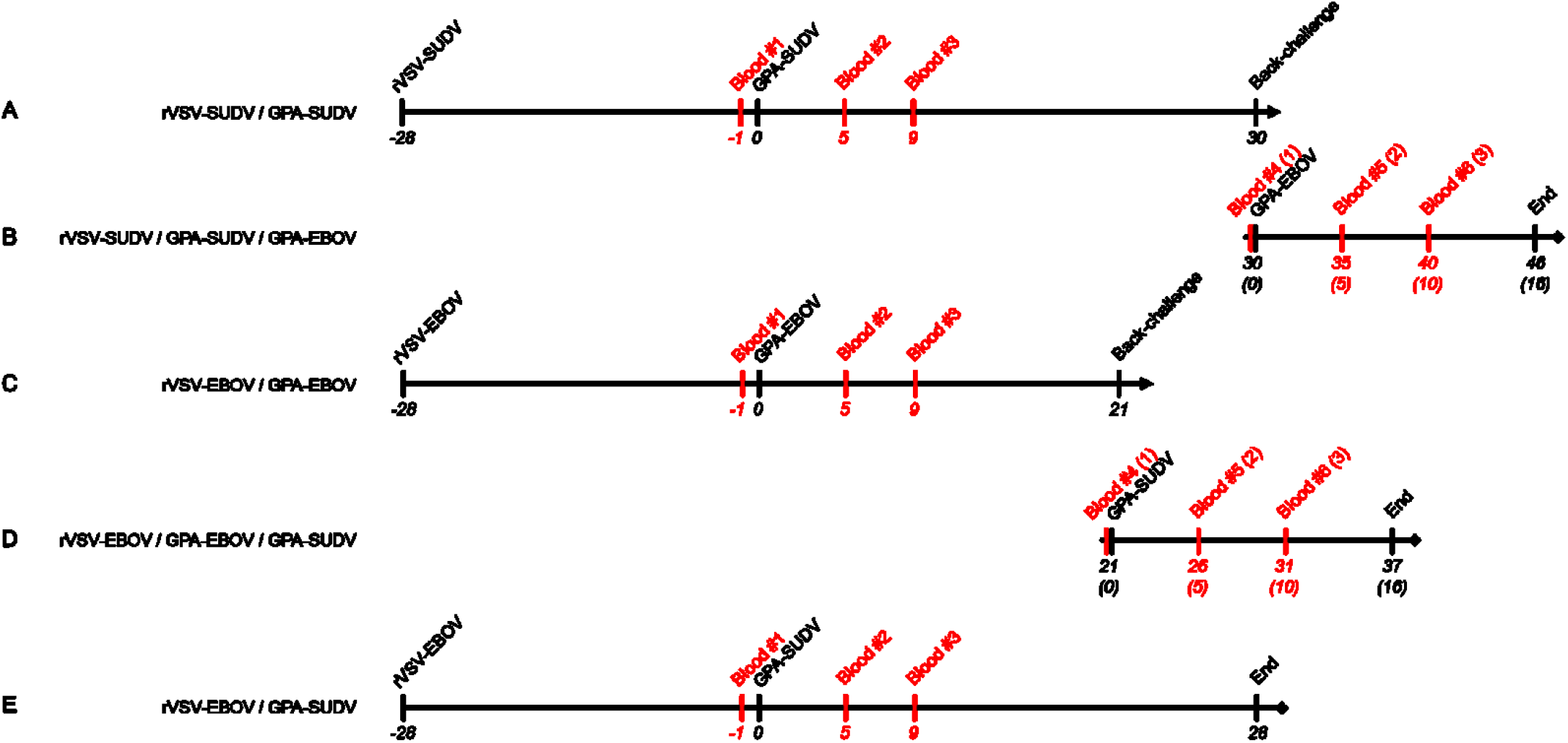
Study Outline. A graphical depiction of the experimental timelines for each challenge experiment. (A) To assess rVSV-SUDV protection against GPA-SUDV challenge, guinea pigs were immunized with 2 × 10^5^ PFU or 2 × 10^3^ PFU rVSV-SUDV and then challenged with 1000 LD_50_ GPA-SUDV 28 days later. (B) On day 30 post-infection, the surviving animals from (A) were back-challenged with 1000 LD_50_ of GPA-EBOV to assess cross-protection. (C) To assess rVSV-EBOV protection against GPA-EBOV challenge, guinea pigs were immunized with 1 × 10^6^ PFU rVSV-EBOV and then challenged with 1000 LD_50_ GPA-EBOV 28 days later. (D) On day 21 post-infection, the surviving animals from (C) were back-challenged with 1000 LD_50_ of GPA-SUDV to assess cross-protection. (E) To assess rVSV-EBOV cross-protection against GPA-SUDV challenge, guinea pigs were immunized with 1 × 10^6^ PFU rVSV-EBOV and then challenged with 1000 LD_50_ GPA-SUDV 28 days later. The experiment was terminated on day 28 post-infection. In general, blood samples were obtained from all animals prior to challenge (Blood #1, day -1) and again on days 5 (Blood #2) and 9 or 10 (Blood #3). Blood samples (Terminal) were also obtained whenever an animal met the humane criteria for euthanasia.

**Fig. S2.**
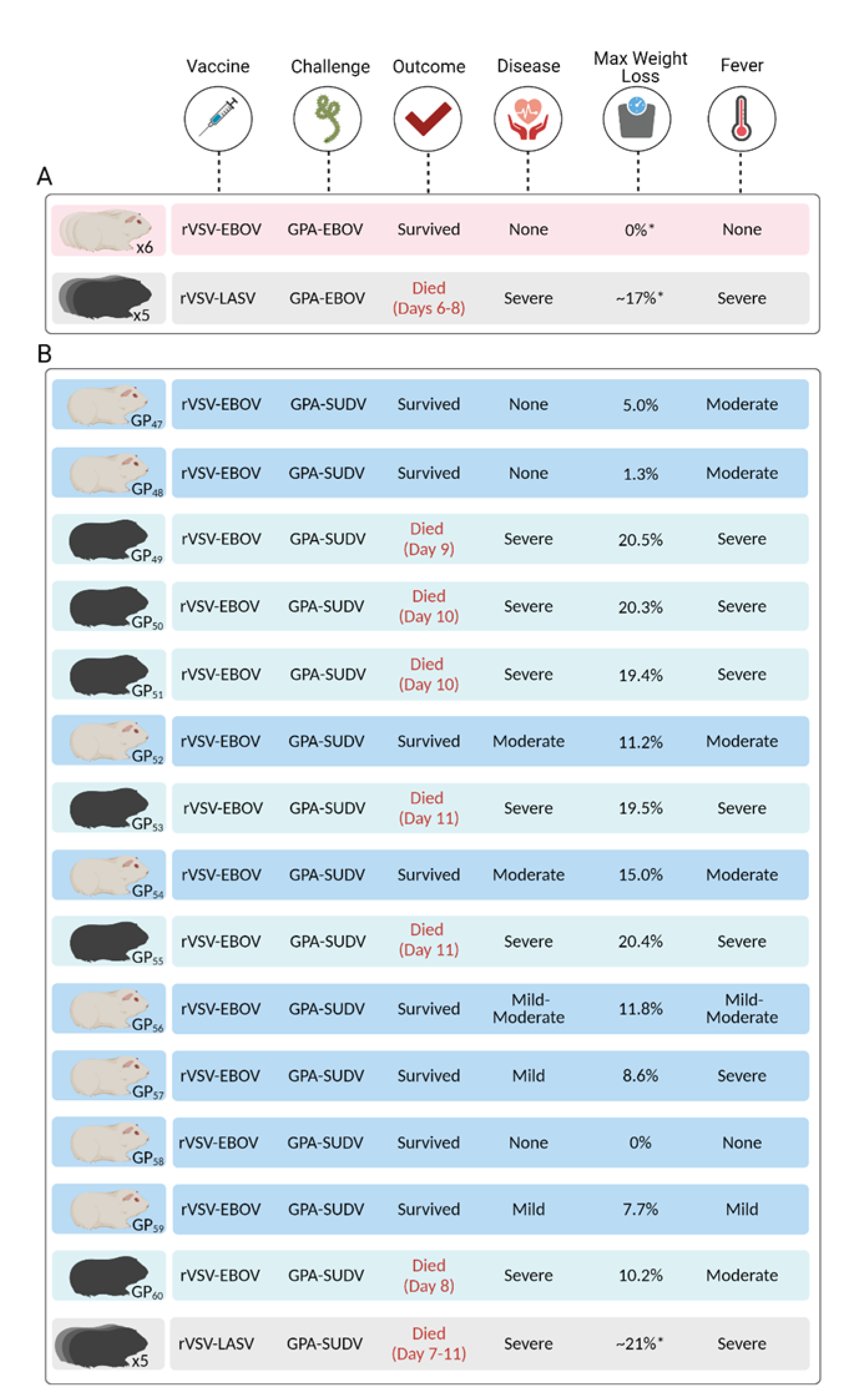
Summary of cross-protection in guinea pigs vaccinated with rVSV-EBOV. The diagram depicts a qualitative summary of the disease observed in guinea pigs vaccinated with rVSV-EBOV or rVSV-LASV and challenged with GPA-EBOV (A) or GPA-SUDV (B). A summary is provided for each of the 14 animals (ID numbers indicated) that were vaccinated with rVSV-EBOV and challenged with GPA-SUDV, while a collective summary is provided for all other animals. Disease severity was qualitatively categorized as severe, moderate, or mild based on the observation of clinical signs, including animal activity and posture, as well as weight loss and body temperature. Fever was categorized based on maximum recorded temperature: >40.5°C, severe; >40°C, moderate; >39.5°C mild. *, indicates an average value. This figure was created with BioRender.com.

**Fig. S3.**
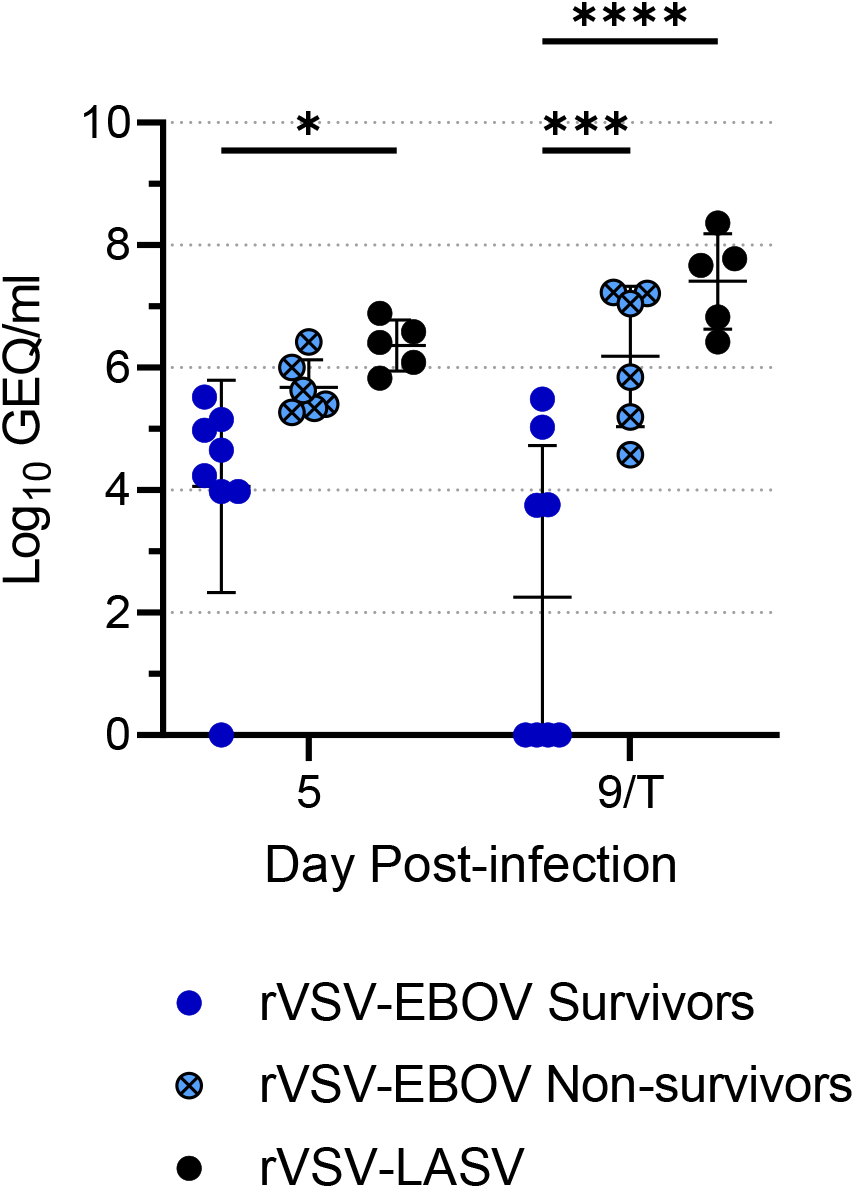
Statistical analysis of virus RNA levels in guinea pigs vaccinated with rVSV-EBOV and challenged with GPA-SUDV. To determine whether the levels of virus RNA were significantly different between the rVSV-EBOV-vaccinated survivors and non-survivors, data from Fig. 4D was re-plotted to separate the rVSV-EBOV vaccinated survivors from the non-survivors. A two-way ANOVA test, along with Tukey’s multiple comparison test, was performed. At day 5, there was no significant difference between the virus RNA levels in rVSV-EBOV vaccinated survivors and non-survivors, although the level of virus RNA in the rVSV-LASV-vaccinated control animals was significantly higher than in the rVSV-EBOV vaccinated survivors. At day 9 or the terminal time point, the average level of virus RNA in the non-survivors was significantly greater than that in the survivors but not significantly different from that in the control animals. Data from animals that were vaccinated with rVSV-EBOV but did not survive challenge with GPA-SUDV are highlighted in light blue and indicated with an “x” on the symbol. *, p ≤ 0.05; ***, p ≤ 0.001; ****, p ≤ 0.0001

**Fig. S4.**
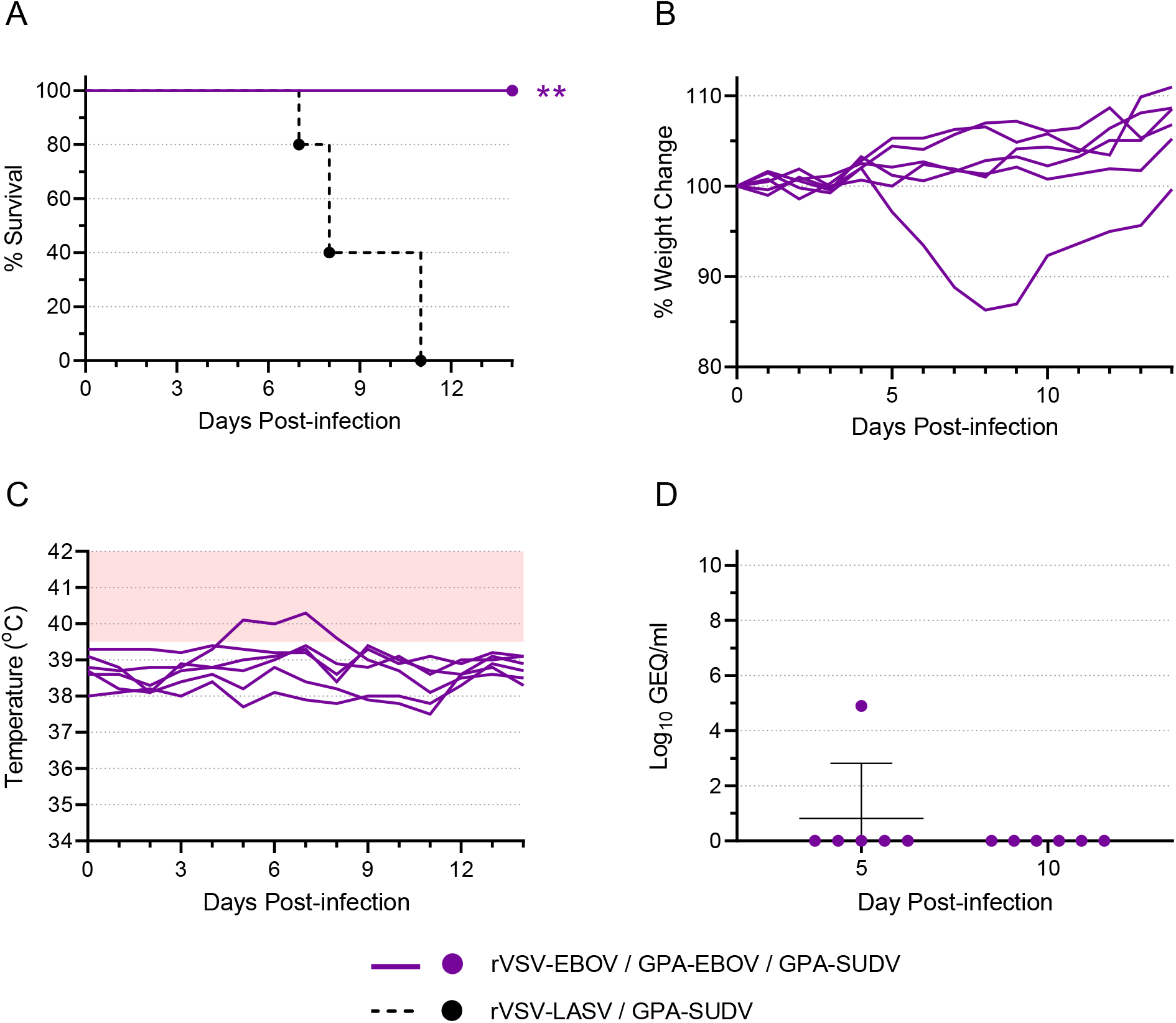
rVSV-EBOV vaccinated survivors from EBOV challenge are protected against SUDV back-challenge. As described for Fig. 3, guinea pigs were vaccinated with rVSV-EBOV (n = 6) and then challenged with 1000 LD_50_ GPA-EBOV. All animals survived GPA-EBOV challenge and, 21 days later, were back-challenged with 1000 LD_50_ of GPA-SUDV. Animals were monitored for survival (A), weight change (B), and body temperature (C). The area shaded pink in (C) highlights temperatures above 39.5 °C. For comparison, the survival curve from animals vaccinated with rVSV-LASV and challenged with GPA-SUDV (Fig. 4) is shown as a dotted line (A). Blood samples were obtained from each animal on day 5 and 10 post-infection. Samples were assessed for levels of virus RNA via RT-qPCR, and data are presented as Log_10_ genome equivalents (GEQ) per milliliter for each animal, with means and standard deviations indicated (D). **, p ≤ 0.01.

**Fig. S5.**
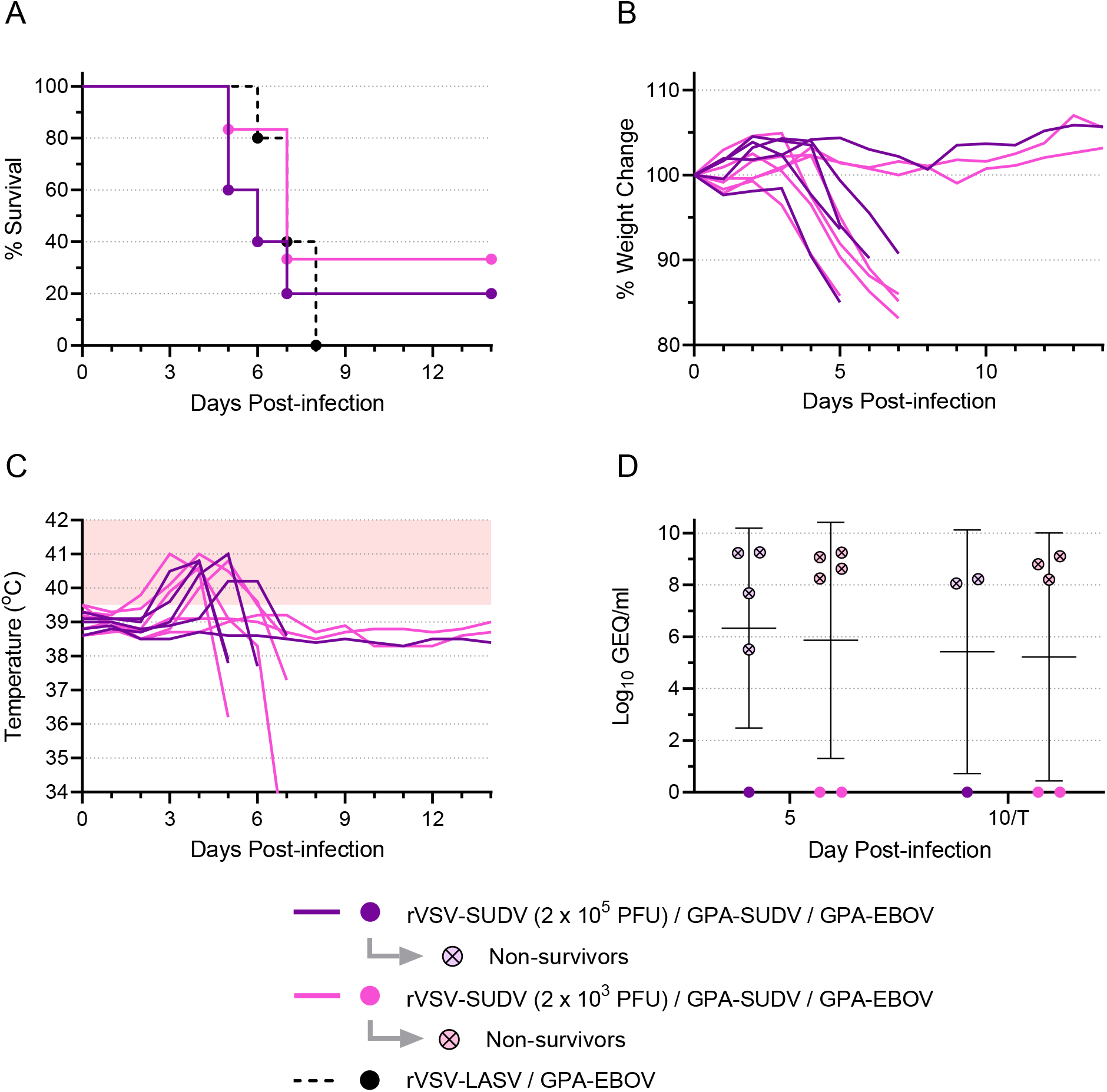
rVSV-SUDV vaccinated survivors from SUDV challenge are not protected against EBOV back-challenge. As described for Fig. 1, guinea pigs were vaccinated with rVSV-SUDV at a dose of either 2 × 10^5^ PFU (n = 6) or 2 × 10^3^ PFU (n = 6) and then challenged with 1000 LD_50_ of GPA-SUDV. All animals survived GPA-SUDV challenge and, 30 days later, were back-challenged with 1000 LD_50_ of GPA-EBOV. Note that a single vaccinated animal died during sampling in the original experiment, so the rVSV-SUDV (2 × 10^5^ PFU) group contained only 5 animals for the back challenge. Animals were monitored for survival (A), weight change (B), and body temperature (C). The area shaded pink in (C) highlights temperatures above 39.5 °C. For comparison, the survival curve from animals vaccinated with rVSV-LASV and challenged with GPA-EBOV (Fig. 3) is shown as a dotted line (A). Blood samples were obtained from each animal on day 5 post-infection and either day 10 post-infection or at the terminal time point (T) if it occurred before day 10. Samples were assessed for levels of virus RNA via RT-qPCR, and data are presented as Log_10_ genome equivalents (GEQ) per milliliter for each animal, with means and standard deviations indicated (D). Data from animals that were back-challenged with GPA-EBOV but did not survive are highlighted in light purple or light pink and indicated with an “x” on the symbol.

## References

1. J. H. Kuhn, G. K. Amarasinghe, D. L. Perry, in Fields Virology: Emerging Viruses, P. M. Howley, D.M. Knipe, S. Whelan, Eds. (Wolters Kluwer, 2021).

2. E. Mahase, Guinea reports west Africa’s first ever case of Marburg virus disease. BMJ 374, 1988 (2021).

3. Y. Araf, S. T. Maliha, J. Zhai, C. Zheng, Marburg virus outbreak in 2022: a public health concern. The Lancet Microbe (2022), doi:10.1016/S2666-5247(22)00258-0.

4. E. I. Ohimain, D. Silas-Olu, The 2013–2016 Ebola virus disease outbreak in West Africa. Curr. Opin. Pharmacol. 60, 360–365 (2021).

5. Ebola outbreak 2018-2020-North Kivu-Ituri (available at https://www.who.int/emergencies/situations/Ebola-2019-drc-).

6. R. Burk, L. Bollinger, J. C. Johnson, J. Wada, S. R. Radoshitzky, G. Palacios, S. Bavari, P. B. Jahrling, J. H. Kuhn, Neglected filoviruses. FEMS Microbiol. Rev. 40, 494–519 (2016).

7. Ebola Virus Disease in Uganda SitRep - 46 | WHO | Regional Office for Africa (available at https://www.afro.who.int/countries/uganda/publication/ebola-virus-disease-uganda-sitrep-46).

8. I. Crozier, K. A. Britson, D. N. Wolfe, J. D. Klena, L. E. Hensley, J. S. Lee, L. A. Wolfraim, K. L. Taylor, E. S. Higgs, J. M. Montgomery, K. A. Martins, The Evolution of Medical Countermeasures for Ebola Virus Disease: Lessons Learned and Next Steps. Vaccines 10 (2022), doi:10.3390/vaccines10081213.

9. M. Garbutt, R. Liebscher, V. Wahl-Jensen, S. Jones, P. Möller, R. Wagner, V. Volchkov, H.-D. Klenk, H. Feldmann, U. Ströher, Properties of Replication-Competent Vesicular Stomatitis Virus Vectors Expressing Glycoproteins of Filoviruses and Arenaviruses. J. Virol. 78, 5458–5465 (2004).

10. E. Ollmann Saphire, A Vaccine against Ebola Virus. Cell 181, 6 (2020).

11. G. Liu, W. Cao, A. Salawudeen, W. Zhu, K. Emeterio, D. Safronetz, L. Banadyga, Vesicular stomatitis virus: From agricultural pathogen to vaccine vector. Pathogens 10, 1092 (2021).

12. Sudan Virus Vaccine Tracker - List of vaccine candidates in research & development (available at https://www.who.int/publications/m/item/sudan-virus-vaccine-tracker---list-of-vaccine-candidates-in-research---development).

13. Ebola Sudan Chimpanzee Adenovirus Vector Vaccine in Healthy Adults - Full Text View - ClinicalTrials.gov (available at https://clinicaltrials.gov/ct2/show/NCT04041570?cond=sudan+ebola&draw=2&rank=3).

14. A Study of a New Vaccine Against Two Types of Ebola - Full Text View - ClinicalTrials.gov (available at https://clinicaltrials.gov/ct2/show/NCT05079750?cond=sudan+ebola&draw=2&rank=1).

15. A Study in Tanzania of a New Vaccine Against Two Types of Ebola - Full Text View - ClinicalTrials.gov (available at https://clinicaltrials.gov/ct2/show/NCT05301504?cond=sudan+ebola&draw=2&rank=2).

16. I. D. Milligan, M. M. Gibani, R. Sewell, E. A. Clutterbuck, D. Campbell, E. Plested, E. Nuthall, M. Voysey, L. Silva-Reyes, M. J. McElrath, S. C. De Rosa, N. Frahm, K. W. Cohen, G. Shukarev, N. Orzabal, W. Van Duijnhoven, C. Truyers, N. Bachmayer, D. Splinter, N. Samy, M. G. Pau, H. Schuitemaker, K. Luhn, B. Callendret, J. Van Hoof, M. Douoguih, K. Ewer, B. Angus, A. J. Pollard, M. D. Snape, Safety and Immunogenicity of Novel Adenovirus Type 26- and Modified Vaccinia Ankara-Vectored Ebola Vaccines: A Randomized Clinical Trial. JAMA 315, 1610–1623 (2016).

17. K. C. L. Shaffer, S. Hui, A. Bratcher, L. B. King, R. Mutombe, N. Kavira, J. P. Kompany, M. Tambu, K. Musene, P. Mukadi, P. Mbala, A. Gadoth, B. R. West, B. K. Ilunga, D. Kaba, J. J. Muyembe-Tanfum, N. A. Hoff, A. W. Rimoin, E. O. Saphire, Pan-ebolavirus serology study of healthcare workers in the Mbandaka Health Region, Democratic Republic of the Congo. PLoS Negl. Trop. Dis. 16 (2022), doi:10.1371/journal.pntd.0010167.

18. A. MacNeil, Z. Reed, P. E. Rollin, Serologic cross-reactivity of human IgM and IgG antibodies to five species of Ebola virus. PLoS Negl. Trop. Dis. 5 (2011), doi:10.1371/journal.pntd.0001175.

19. M. Natesan, S. M. Jensen, S. L. Keasey, T. Kamata, A. I. Kuehne, S. W. Stonier, J. J. Lutwama, L. Lobel, J. M. Dye, R. G. Ulrich, Human survivors of disease outbreaks caused by Ebola or Marburg virus exhibit cross-reactive and long-lived antibody responses. Clin. Vaccine Immunol. 23, 717–724 (2016).

20. X. Yu, E. O. Saphire, Development and Structural Analysis of Antibody Therapeutics for Filoviruses. Pathogens 11, 374 (2022).

21. S. M. Jones, H. Feldmann, U. Ströher, J. B. Geisbert, L. Fernando, A. Grolla, H. D. Klenk, N. J. Sullivan, V. E. Volchkov, E. A. Fritz, K. M. Daddario, L. E. Hensley, P. B. Jahrling, T. W. Geisbert, Live attenuated recombinant vaccine protects nonhuman primates against Ebola and Marburg viruses. Nat. Med. 11, 786–790 (2005).

22. T. W. Geisbert, J. B. Geisbert, A. Leung, K. M. Daddario-DiCaprio, L. E. Hensley, A. Grolla, H. Feldmann, Single-Injection Vaccine Protects Nonhuman Primates against Infection with Marburg Virus and Three Species of Ebola Virus. J. Virol. 83, 7296–7304 (2009).

23. A. Marzi, F. Engelmann, F. Feldmann, K. Haberthur, W. L. Shupert, D. Brining, D. P. Scott, T. W. Geisbert, Y. Kawaoka, M. G. Katze, H. Feldmann, I. Messaoudi, Antibodies are necessary for rVSV/ZEBOV-GP-mediated protection against lethal Ebola virus challenge in nonhuman primates. Proc. Natl. Acad. Sci. U. S. A. 110, 1893–1898 (2013).

24. L. Banadyga, G. Wong, X. Qiu, Small Animal Models for Evaluating Filovirus Countermeasures. ACS Infect. Dis. 4, 673–685 (2018).

25. A. C. Shurtleff, T. K. Warren, S. Bavari, Nonhuman primates as models for the discovery and development of ebolavirus therapeutics. Expert Opin. Drug Discov. 6, 233–250 (2011).

26. A. Marzi, H. Ebihara, J. Callison, A. Groseth, K. J. Williams, T. W. Geisbert, H. Feldmann, Vesicular stomatitis virus-based Ebola vaccines with improved cross-protective efficacy. J. Infect. Dis. 204 (2011), doi:10.1093/infdis/jir348.

27. D. Falzarano, F. Feldmann, A. Grolla, A. Leung, H. Ebihara, J. E. Strong, A. Marzi, A. Takada, S. Jones, J. Gren, J. Geisbert, S. M. Jones, T. W. Geisbert, H. Feldmann, Single immunization with a monovalent vesicular stomatitis virus-based vaccine protects nonhuman primates against heterologous challenge with bundibugyo ebolavirus. J. Infect. Dis. 204 (2011), doi:10.1093/infdis/jir350.

28. C. E. Mire, J. B. Geisbert, A. Marzi, K. N. Agans, H. Feldmann, T. W. Geisbert, Vesicular Stomatitis Virus-Based Vaccines Protect Nonhuman Primates against Bundibugyo ebolavirus. PLoS Negl. Trop. Dis. 7 (2013), doi:10.1371/journal.pntd.0002600.

29. A. Marzi, P. Fletcher, F. Feldmann, G. Saturday, P. W. Hanley, H. Feldmann, VSV-based vaccine provides species-specific protection against Sudan virus challenge in macaques. bioRxiv, 2022.10.27.514045 (2022).

30. R. W. Cross, K. A. Fenton, T. W. Geisbert, Small animal models of filovirus disease: recent advances and future directions. Expert Opin. Drug Discov. 13, 1027–1040 (2018).

31. L. Banadyga, M. A. Dolan, H. Ebihara, Rodent-Adapted Filoviruses and the Molecular Basis of Pathogenesis. J. Mol. Biol. 428, 3449–3466 (2016).

32. G. Wong, S. He, H. Wei, A. Kroeker, J. Audet, A. Leung, T. Cutts, J. Graham, D. Kobasa, C. Embury-Hyatt, G. P. Kobinger, X. Qiu, Development and Characterization of a Guinea Pig-Adapted Sudan Virus. J. Virol. 90, 392–399 (2015).

33. T. W. Geisbert, K. M. Daddario-DiCaprio, K. J. N. Williams, J. B. Geisbert, A. Leung, F. Feldmann, L. E. Hensley, H. Feldmann, S. M. Jones, Recombinant Vesicular Stomatitis Virus Vector Mediates Postexposure Protection against Sudan Ebola Hemorrhagic Fever in Nonhuman Primates. J. Virol. 82, 5664–5668 (2008).

34. B. M. Connolly, K. E. Steele, K. J. Davis, T. W. Geisbert, W. M. Kell, N. K. Jaax, P. B. Jahrling, Pathogenesis of experimental Ebola virus infection in guinea pigs. J. Infect. Dis. 179 (1999), doi:10.1086/514305.

